# Directed effective connectivity and synaptic weights of in vitro neuronal cultures revealed from high-density multielectrode array recordings

**DOI:** 10.1101/2020.02.06.936781

**Authors:** Chumin Sun, K.C. Lin, Yu-Ting Huang, Emily S.C. Ching, Pik-Yin Lai, C.K. Chan

## Abstract

Studying connectivity of neuronal cultures can provide insights for understanding brain networks but it is challenging to reveal neuronal connectivity from measurements. We apply a novel method that uses a theoretical relation between the time-lagged cross-covariance and the equal-time cross-covariance to reveal directed effective connectivity and synaptic weights of cortical neuron cultures at different days in vitro from multielectrode array recordings. Using a stochastic leaky-integrate-and-fire model, we show that the simulated spiking activity of the reconstructed networks can well capture the measured network bursts. The neuronal networks are found to be highly nonrandom with an over-representation of bidirectionally connections as compared to a random network of the same connection probability, with the fraction of inhibitory nodes comparable to the measured fractions of inhibitory neurons in various cortical regions in monkey, and have small-world topology with basic network measures comparable to those of the nematode C. elegans chemical synaptic network. Our analyses further reveal that (i) the excitatory and inhibitory incoming degrees have bimodal distributions the excitatory and inhibitory incoming degrees have bimodal distributions, which are that distributions that have been indicated to be optimal against both random failures and attacks in undirected networks; (ii) the distribution of the physical length of excitatory incoming links has two peaks indicating that excitatory signal is transmitted at two spatial scales, one localized to nearest nodes and the other spatially extended to nodes millimeters away, and the shortest links are mostly excitatory towards excitatory nodes and have larger synaptic weights on average; (iii) the average incoming and outgoing synaptic strength is non-Gaussian with long tails and, in particular, the distribution of outgoing synaptic strength of excitatory nodes with excitatory incoming synaptic strength is lognormal, similar to the measured excitatory postsynaptic potential in rat cortex.

**Author summary:** To understand how the brain processes signal and carries out its function, it is useful to know the connectivity of the underlying neuronal circuits. For large-scale neuronal networks, it is difficult to measure connectivity directly using electron microscopy techniques and methods that can estimate connectivity from electrophysiological recordings are thus highly desirable. Existing methods focus mainly on estimating functional connectivity, which is defined by statistical dependencies between neuronal activities but the relevant direct casual interactions are captured by effective connectivity. Here we apply a novel covariance-relation based method to estimate the directed effective connectivity and synaptic weights of cortical neuron cultures from recordings of multielectrode array of over 4000 electrodes taken at different days in vitro. The neuronal networks are found to be nonrandom, small-world, excitation/inhibition balanced as measured in monkey cortex, and with feeder hubs. Our analyses further suggest some form of specialisation of nodes in receiving excitatory and inhibitory signals and the transmission of excitatory signals at two spatial scales, one localized to nearest nodes and the other spatially extended to nodes millimeters away, and reveal that the distributions of the average incoming and outgoing synaptic strength are skewed with long tails.

## Introduction

Unravelling synaptic connectivity and understanding the relationship between connectivity and dynamics and between network structure and function of neuronal circuits are actively pursued topics in neuroscience [1, 2]. In vitro neuronal cultures serve as simple yet useful experimental model systems for studying the relationship between connectivity and the rich dynamics observed [3–5]. One common technique used to record the activity of neurons in cultures is direct measurement of the electrical signals of neurons using multielectrode array (MEA) [6]. It is thus of great interest to estimate connectivity from MEA recordings. There is a distinction between functional and effective connectivity [7]. Functional connectivity is defined by statistical dependencies between the activities of neurons and is usually inferred based on the cross-covariance or cross-correlation of spike trains with the spikes first being detected from the recorded electrical signals of the electrodes of the MEA [8–10]. As statistical dependencies can arise from indirect interactions, functional connectivity does not necessarily reflect the direct causal interactions among the neurons. Direct causal interaction is referred to as effective connectivity, and is more relevant for studying the relationship between connectivity and dynamics and between network structure and function. Effective connectivity is used to be inferred by using some specific model of interactions and, as a result, it is sometimes said to be model-dependent. Reconstructing functional connectivity from measurements is already highly challenging and nontrivial especially for large-scale MEA with several thousands of electrodes. Existing methods thus focus predominantly on estimating functional connectivity.

For a class of networked systems having generic stationary dynamics modelled by a set of stochastic differential equations with directional interactions, it has been shown that the equal-time cross-covariance of the dynamics of the nodes does not carry sufficient information to recover the effective connectivity [11, 12], and the effective connectivity matrix can instead be extracted using the theoretical relation between the time-lagged cross-covariance and the equal-time cross-covariance [13]. Similar result has been found for systems with discrete-time dynamics [14]. This covariance-relation based method thus infers effective connectivity for a generic class of models and not only for a specific model. It has been validated by numerical simulations for weighted directed random and scale-free networks with different nonlinear dynamics and directional coupling including particularly the FitzHugh-Nagamo dynamics [15] with directional synaptic-like coupling [13, 16].

In this paper, we show how we adopt this covariance-relation based method to estimate directed effective connectivity of cultures of cortices of rat embryos from recordings of large-scale MEA of over 4000 electrodes taken at different times, ranging from 11 to 66 days in vitro. This method makes use of the MEA recordings directly without the need to first detect spikes from the recordings. In addition, it can detect unidirectional and bidirectional connections equally well and further allows one to estimate the synaptic weights of the links, both excitatory and inhibitory. For cross-correlation based methods that estimate functional connectivity, inhibitory links pose additional challenges [10] and bidirectional connections are difficult to detect. As the validity of the covariance-relation based method has already been shown using numerical data, we focused to test how well the estimated directed effective connectivity can reproduce the measured spiking activity using a stochastic leaky-integrate-and-fire model. We carried out analyses of the reconstructed networks studying the basic network features and the distributions of incoming and outgoing degrees as well as the average incoming and outgoing synaptic strength.

## Results

Cortices from embryonic day 17 (E17) Wistar rats were dissected and a total of about 6 × 10^4^ cells were plated on a 6mm by 6mm working area of a high-density MEA probe. The MEA probe has 4096 electrodes arranged on a 64 by 64 square grid in the central 2.67mm by 2.67mm active area. We recorded spontaneous activity at a sampling frequency of 7.06kHz at eight different Days In Vitro (DIV): 11, 22, 25, 33, 45, 52, 59 and 66. The electrode in the upper left corner was reserved for calibration purpose so 4095 electrodes recorded data. As in earlier work [8], we treated each electrode as a node of the neuronal network and estimated the 4095 × 4095 directed and weighted effective connectivity matrix for all 8 different DIVs, referred to as DIV11, DIV22, DIV25, DIV33, DIV45, DIV52, DIV59 and DIV66, from the relation between the time-lagged covariance and equal-time covariance (c.f. Materials and methods). The validity of this covariance-relation based method in recovering the directed and weighted effective connectivity for random and scale-free networks with a generic model of dynamics has been verified using numerical simulations [13, 16]. In this work, we tested how well the estimated directed and weighted effective connectivity matrix can reproduce the measured spiking activity. Then we analysed the basic features of the reconstructed neuronal networks and calculated the distributions of degree and synaptic strength.

### Testing how well the estimated directed effective connectivity can reproduce the measured spiking activity

We applied the Precision Timing Spike Detection (PTSD) algorithm [17] to the MEA recordings to obtain the measured spiking activity. We used the stochastic leaky integrate-and-fire (LIF) model (c.f. Materials and methods for the computer model) of 4095 neurons coupled according to the reconstructed directed and weighted effective connectivity matrix to simulate the spiking activity. In the LIF model, which is a simple, intuitive and most widely used spiking neuron model [18], the membrane potential for each neuron changes in time due to the inputs from its pre-synaptic neurons and when it reaches a threshold, the neuron generates an action potential or a spike and the membrane potential is reset to the resting state. We added a noise to the usual LIF model [19] acting on each neuron to drive the spontaneous spiking activity of interest. For each neuron, we simulated the spiking activity using the measured spiking activity of all its pre-synaptic neurons as inputs. We carried out the simulations for DIV25, DIV45, and DIV66 for a time window of observation *T*_*obs*_ ranging between 18.9s to 35.9s. In Fig 1, we compared the raster plots of the simulated spiking activity with the measured spiking activity. For a close-up comparison, the plots focus on the 384 nodes corresponding to the electrodes in the 6 middle rows of the 64 by 64 MEA. An outstanding feature of the measured spiking activity is the collective bursts of activity by almost the entire network. The duration of these network bursts decreases as the age of the cultures increases from 25 to 66 DIV. The simulated spiking activity of the reconstructed networks captures such network bursts well.

**Fig 1.**
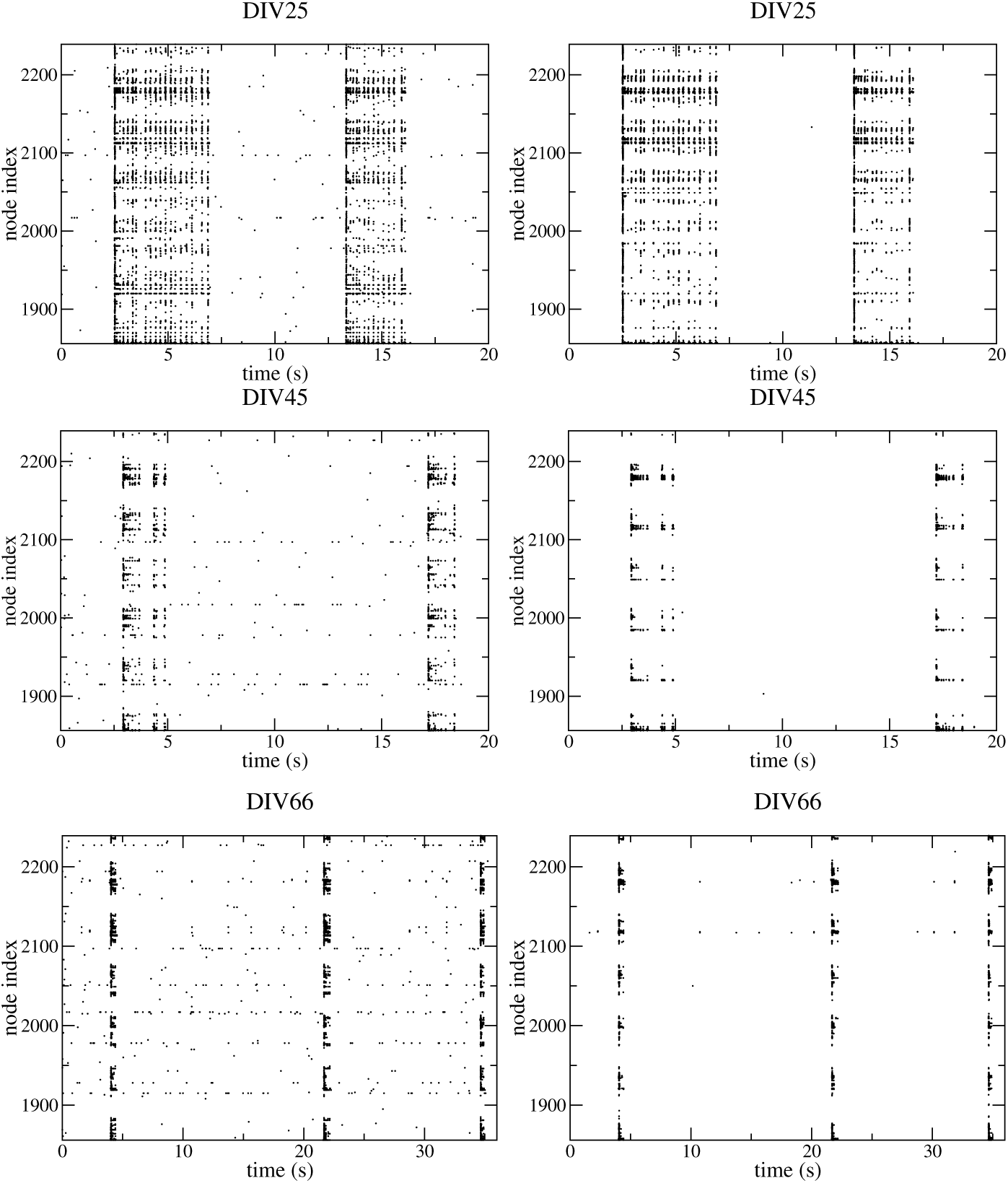
Comparison of the measured spiking activity (left panel) with the simulated spiking activity using the reconstructed network (right panel) for DIV25, DIV45 and DIV66.

A significant fraction of nodes participated in the network bursts but some nodes did not. We denoted nodes that have no measured spiking activity in *T*_*obs*_ as null-nodes and checked how well the simulated results managed to retrieve these null-nodes. As a control, we randomly shuffled the reconstructed network with the distributions of the incoming and outgoing degrees kept intact and assigned the synaptic weights randomly from a Gaussian distribution of zero mean and unit variance and repeated the simulations. We compared how well the simulated results from the reconstructed networks and the control networks managed to retrieve the null-nodes by calculating the sensitivity, which measures the fraction of null-nodes that is correctly retrieved, and the precision, which measures the fraction of nodes simulated to be null-nodes are indeed measured null-nodes. Denote the number of null-nodes that have null or nonzero simulated spiking activity by *N*_*TP*_ or *N*_*FN*_ and the number of non null-nodes with null or nonzero simulated spiking activity by *N*_*FP*_ or *N*_*TN*_. The sensitivity (SEN) and precision, which is also known as the positive predictive value (PPV) are defined by

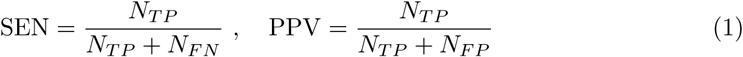

The results obtained are presented in Table 1. Both SEN and PPV are much higher for the reconstructed networks than for the control randomly-shuffled networks. Thus the reconstructed networks outperform the control randomly reshuffled networks in retrieving the null nodes.

**Table 1.** The sensitivity and precision or positive predictive value of the reconstructed and control randomly-shuffled networks in retrieving null-nodes. *f* is the fraction of the *N*_*TN*_ nodes that have correlation coefficient *ρ* ≥ 0.8.

| DIV | Network | $N_{TP}$ | $N_{FN}$ | $N_{FP}$ | $N_{TN}$ | SEN | PPV | $f$ |
| --- | --- | --- | --- | --- | --- | --- | --- | --- |
| 25 | reconstructed | 222 | 370 | 236 | 3267 | 0.38 | 0.48 | 0.32 |
|  | control | 39 | 553 | 146 | 3357 | 0.066 | 0.21 | 0.059 |
| 45 | reconstructed | 1171 | 453 | 755 | 1716 | 0.72 | 0.61 | 0.47 |
|  | control | 347 | 1277 | 297 | 2174 | 0.21 | 0.54 | 0.14 |
| 66 | reconstructed | 781 | 490 | 656 | 2168 | 0.61 | 0.54 | 0.58 |
|  | control | 97 | 1174 | 119 | 2705 | 0.076 | 0.45 | 0.35 |

We further studied the correlation of the measured and simulated spiking activity at tens of millisecond timescale. We calculated the Pearson correlation coefficient *ρ* between the measured and simulated spiking activity using a sliding window of *w* = 28.4 ms for each of the *N*_*TN*_ nodes with non-zero measured and simulated spiking activity. The distributions of the correlation coefficient *P*(*ρ*) are shown in Fig 2. The fraction *f* of these *N*_*TN*_ nodes with *ρ* ≥ 0.8 is significantly larger for the simulated results using the reconstructed networks (see Table 1) than those using the control randomly shuffled networks.

**Fig 2.**
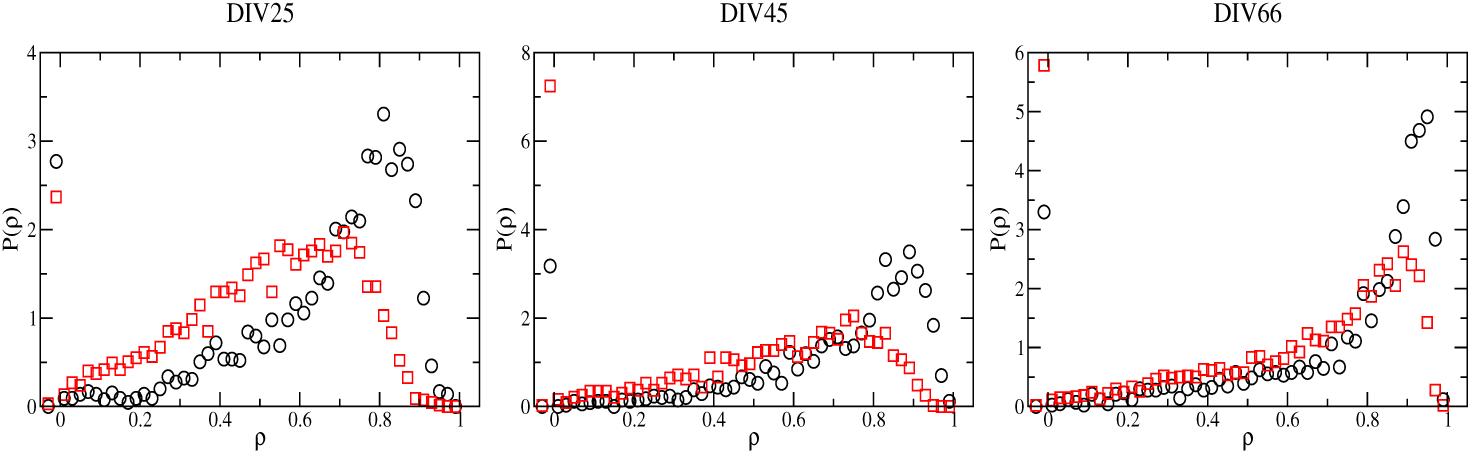
Distributions of the correlation coefficient *P* (*ρ*) for all the *N*_*TN*_ nodes for DIV25, DIV45 and DIV66. Black circles are results for the simulations using the reconstructed network and red squares are results for simulations using the control randomly-shuffled network.

All these results further support the validity of the covariance-relation based method in estimating directed effective connectivity. In the next subsections, we show results of the basic features and the distributions of degrees and synaptic strength of the reconstructed neuronal networks.

### Basic network features

We calculated several basic features of the neuronal networks for all the 8 DIVs (see Table 2). The connection probability *p* of a network of *N* nodes with *N*_*L*_ links is defined by *p* = *N*_*L*_*/*[*N*(*N* − 1)]. We found that *p* ranges from 1.1-1.9%, confirming that the neuronal networks are sparse. This value of *p* is comparable to that of the chemical synapse network of C. elegans [20]. Most of the connections are unidirectional as expected. For a random network of the same number of nodes and the same connection probability *p*, the expected number of bidirectionally connected pairs is given by *N*(*N* − 1)*p*^2^*/*2. The number of bidirectionally connected pairs is indeed comparable to this expected number for the control randomly shuffled networks but exceeds four times this expected number for the reconstructed networks. This observation of an over-representation of bidirectional connections as compared to a random network in the reconstructed neuronal networks echoes previous reports for local regions of the rat cortex [21–24] Thus neuronal networks are highly nonrandom. As a comparison, we calculated the number of bidirectional connections in the functional connectivity estimated by using the cross-covariance based methods [10] for DIV25, DIV45 and DIV66 and found none.

**Table 2.** Basic network measures of the reconstructed networks for the 8 DIVs including the connection probability *p*, the ratio *r*_*B*_ of the number of bidirectionally connected pairs to the expected number *N* (*N* − 1)*p*^2^*/*2 for a random network with *p*, the fractions *f*_*E*_ and *f*_*I*_ of excitatory and inhibitory nodes, the fraction *f*_*SCC*_ of nodes that form the strongly connected component, the characteristic path length *l*, the average clustering coefficient CC and the small world index SWI. When available, the corresponding results for the chemical synapse network of C. elegans [20] are included for comparison.

|  | DIV11 | DIV22 | DIV25 | DIV33 | DIV45 | DIV52 | DIV59 | DIV66 | C. elegans |
| --- | --- | --- | --- | --- | --- | --- | --- | --- | --- |
| $p$ (%) | 1.2 | 1.9 | 1.4 | 1.5 | 1.1 | 1.7 | 1.4 | 1.5 | 2.8 |
| $r_B$ | 5.9 | 5.5 | 10.7 | 5.7 | 6.5 | 4.21 | 4.0 | 4.4 | — |
| $f_E$ | 0.62 | 0.80 | 0.84 | 0.75 | 0.62 | 0.66 | 0.48 | 0.57 | — |
| $f_I$ | 0.27 | 0.13 | 0.14 | 0.18 | 0.21 | 0.24 | 0.32 | 0.28 | — |
| $f_{SCC}$ | 0.88 | 0.93 | 0.98 | 0.92 | 0.81 | 0.90 | 0.77 | 0.83 | 0.85 |
| $l$ | 4.0 | 3.7 | 3.7 | 3.9 | 4.1 | 3.7 | 3.8 | 3.8 | 3.5 |
| CC | 0.26 | 0.36 | 0.38 | 0.30 | 0.25 | 0.28 | 0.18 | 0.22 | 0.22 |
| SWI | 13.1 | 11.3 | 17.1 | 12.7 | 14.4 | 10.4 | 7.9 | 8.8 | 2.3 |

Each node in the neuronal network is inferred to be excitatory or inhibitory according to the sign of the synaptic weights of the outgoing links or has no detectable outgoing links. The spatial distribution of the excitatory and inhibitory nodes and nodes with no detectable outgoing links are shown in Fig 3 for several DIVs. As shown in Table 2, the fraction *f*_*E*_ of excitatory nodes first increases as DIV increases and peaks at DIV25 then generally decreases as DIV further increases while the fraction *f*_*I*_ of inhibitory nodes first decreases and reaches a minimum at DIV22 and then generally increases as DIV further increases. Moreover, *f*_*I*_ ranges from 0.13 to 0.31, which is comparable to the measured fractions (0.15-0.30) of inhibitory neurons in various cortical regions in monkey [25]. The fraction *f*_*SCC*_ of nodes that form the largest strongly connected component exceeds 70% for all the 8 DIVs. The characteristic path length *l*, which is the average shortest path length for each pair of nodes in the strongly connected component, is about 4, and the average local clustering coefficient (CC) measuring the average connection probability for nodes with different outgoing degree ranges from 0.18 − 0.38. These values are comparable to those of the chemical synapse network of C. elegans. Furthermore, the neuronal networks at all the 8 DIVs have small-world topology, with small-world index (SWI) substantially greater than one.

**Fig 3.**
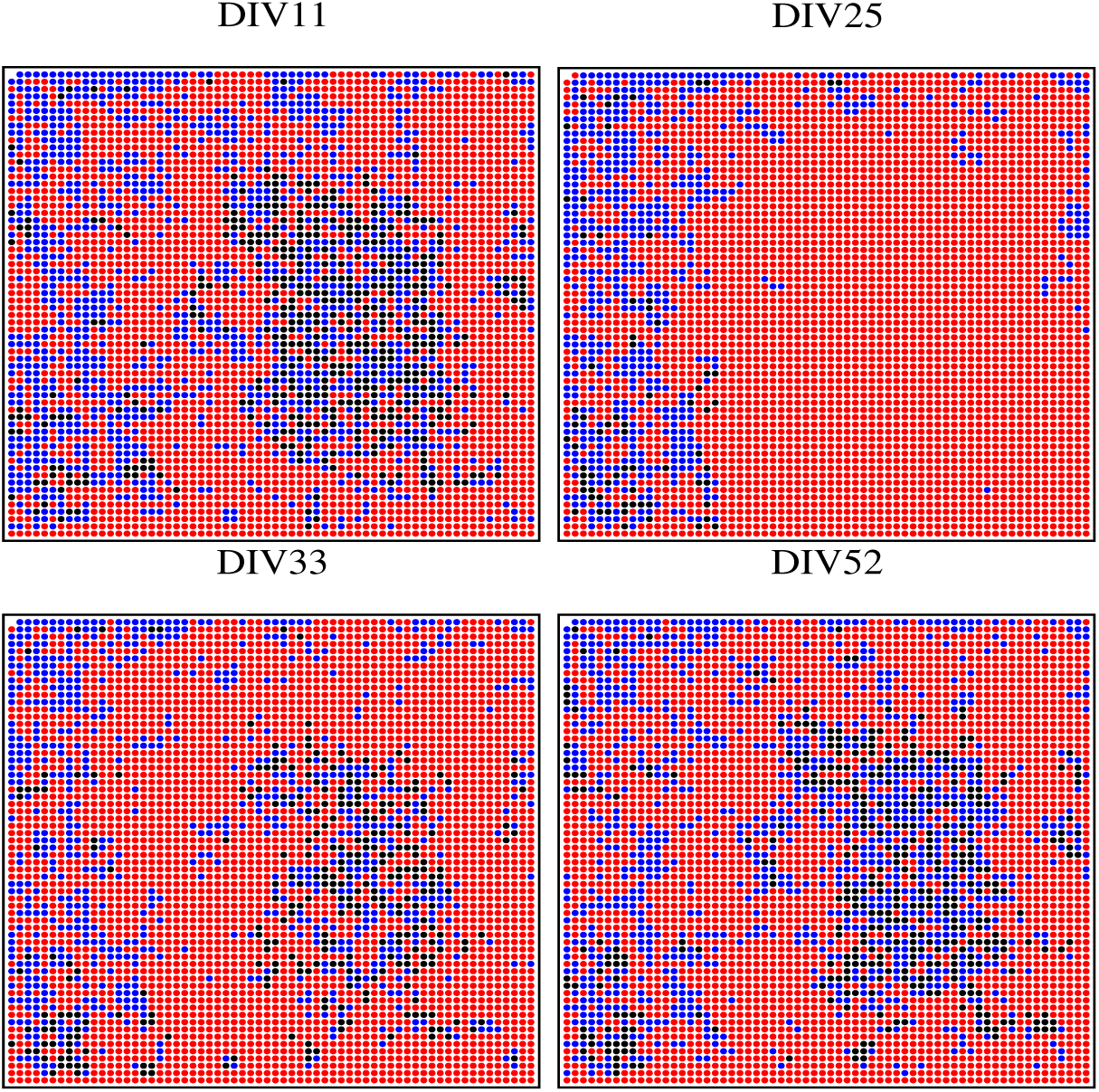
Spatial distribution of excitatory nodes (red), inhibitory nodes (blue) and nodes with no detectable outgoing links (black) for DIV11, DIV25, DIV33 and DIV52.

### Distributions of incoming and outgoing degrees

We calculated both the incoming degree *k*_*in*_ and the outgoing degree *k*_*out*_ for each node, and *k*_*in*_ ≠ *k*_*out*_ as the neuronal networks are directed. The distributions of the incoming and outgoing degrees are qualitatively the same for all the 8 DIVs and the results for DIV25 are shown in Fig 4. A striking feature of the distribution of the incoming degree is its approximate bimodal feature, which is in contrast to the degree distributions of the chemical synapse network of C. elegans [20] and the scale-free distribution found in the functional connectivity [10]. For undirected networks, there have been studies indicating that the robustness against both random failures and attacks can be optimized by having a bimodal degree distribution [26, 27].

**Fig 4.**
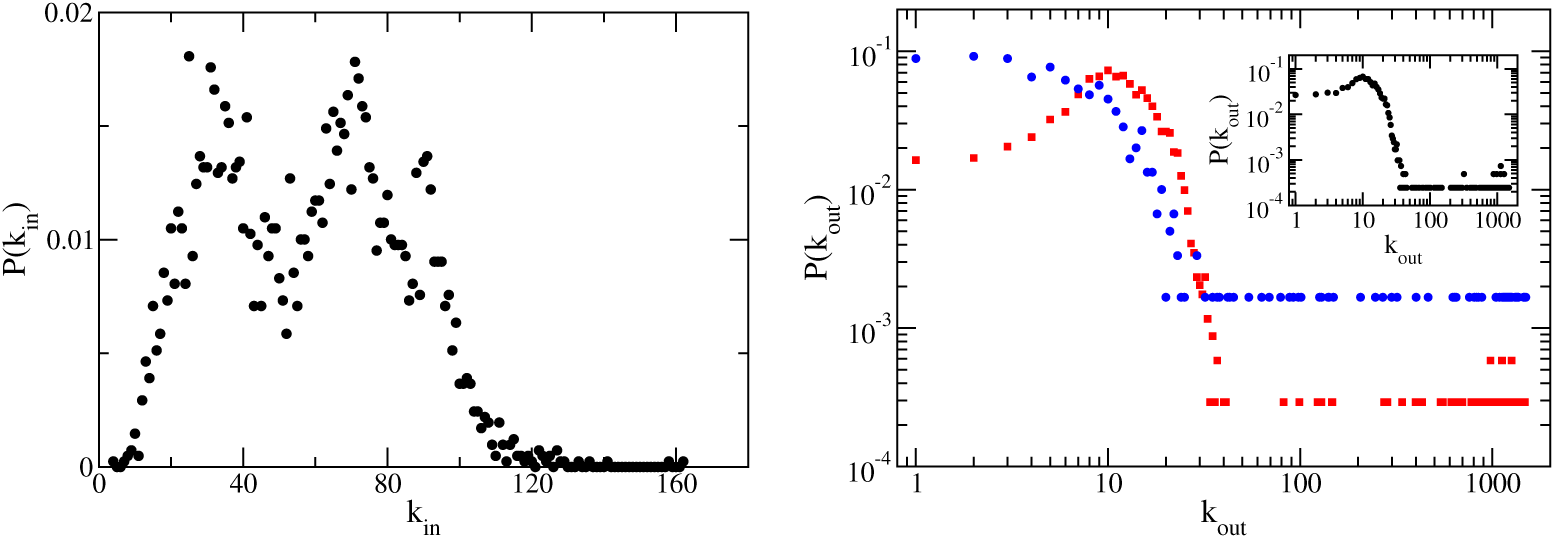
Left panel: Distribution *P* (*k*_*in*_) of the incoming and outgoing degrees for DIV25. Right panel: Distribution *P* (*k*_*out*_) of the outgoing degrees for excitatory (red) and inhibitory nodes (blue) for DIV25. In the inset, we show the overall distribution of *k*_*out*_ for all the nodes.

For DIV25, the outgoing degree of most of the nodes is below 30, which is smaller than the average incoming degree, and a small fraction of nodes have exceptionally large *k*_*out*_ (*k*_*out*_ > 1000). This is the case for both excitatory and inhibitory nodes. Thus the hubs of the neuronal networks are excitatory and inhibitory feeders. The spatial location of these feeder hubs for DIV25 and DIV33 is shown in Fig 5. Interestingly, the feeder hubs locate mostly on the top and bottom rows of the MEA probe.

**Fig 5.**
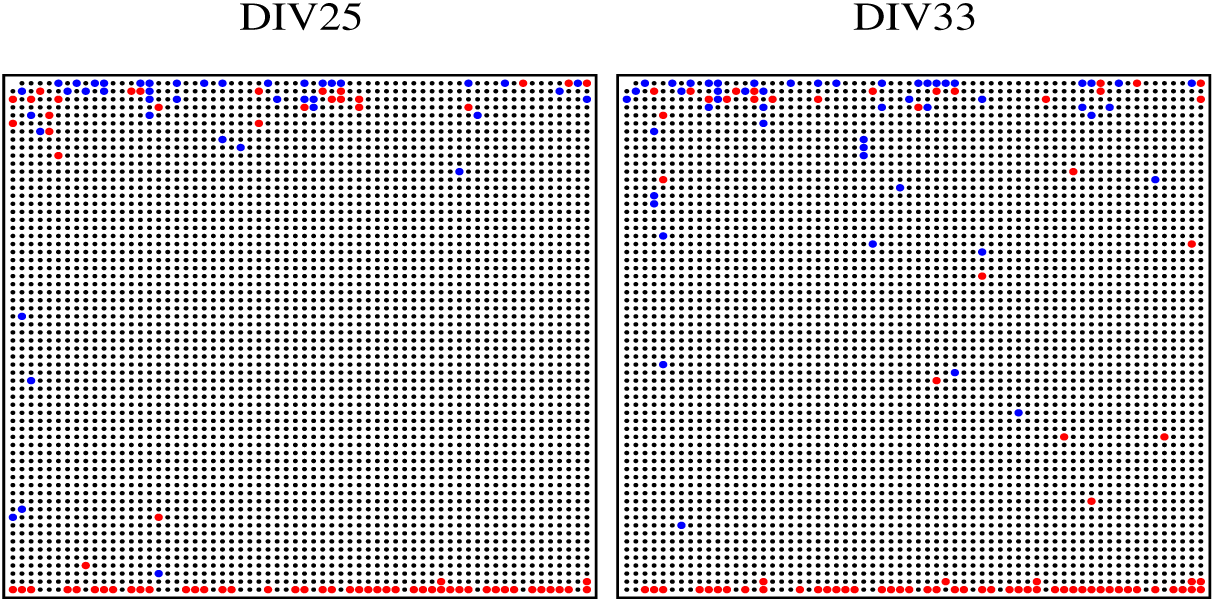
Spatial distribution of excitatory (red) and inhibitory (blue) feeder hubs with *k*_*out*_ > 1000 for DIV25 and DIV33.

To investigate further the approximate bimodal feature of *P* (*k*_*in*_), we calculated separately the distributions of excitatory and inhibitory incoming degrees 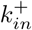 and 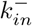 with incoming links of positive and negative *w*_*ij*_ respectively. As shown in Fig 6, both 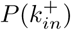 and 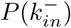 are clearly bimodal and have a smaller mode of larger 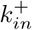 or 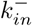. Interestingly, a much reduced fraction of nodes in the mode of larger 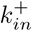 belong to the mode of larger 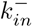, and similarly a much reduced fraction of nodes in the mode of larger 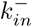 belong to the mode of larger 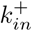 showing that nodes avoid to exhibit both large excitatory and large inhibitory incoming degrees. We show the spatial distribution of nodes in the two modes of the two distributions (see Fig 7). Nodes in the mode of larger 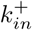 occupy mostly the upper-quarter region while nodes in the mode of larger 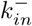 occupy mostly the lower-quarter region showing clearly that nodes avoid having both large excitatory and large inhibitory incoming degrees. This suggests some form of specialization of nodes in receiving excitatory and inhibitory signals.

**Fig 6.**
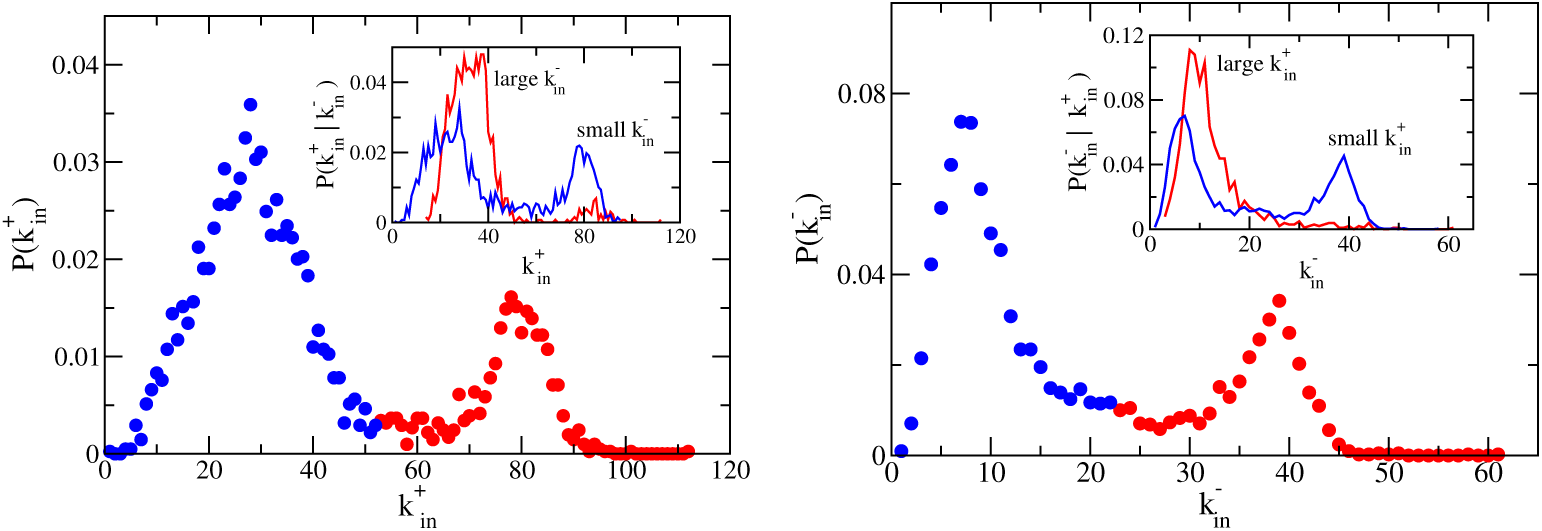
Distributions 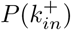 and 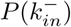 of excitatory and inhibitory incoming degrees for DIV25. The distributions are bimodal with a smaller mode of larger 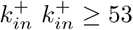) or larger 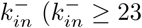 (red) and a larger mode of smaller 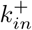 or smaller 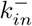 (blue). In the insets, we show the conditional distributions of 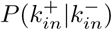 and 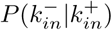 for nodes in the two modes.

**Fig 7.**
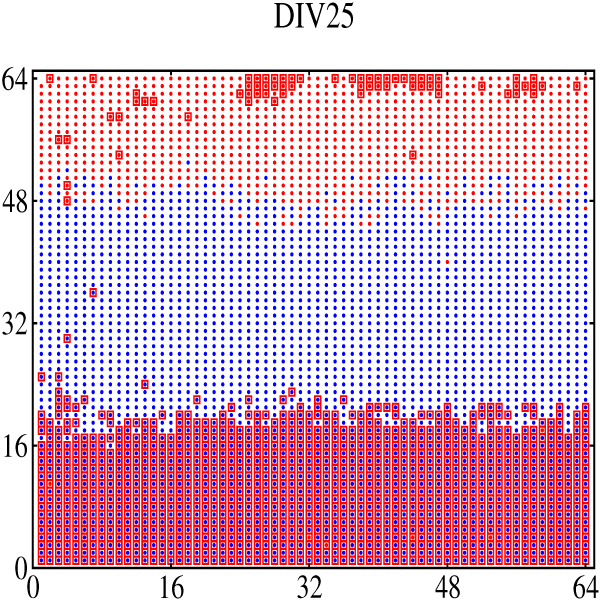
Spatial distribution of nodes in the two modes of larger 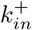 (solid red circles) and smaller 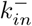 (solid blue circles) and in the two modes of larger 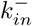 (open red squares) and smaller 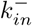 (all symbols without open red squares) for DIV25.

We show the distributions of the physical length of excitatory and inhibitory links in Fig 8. Both distributions have a broad peak at *L* ≈ 60*d* = 2.5mm, where *d* = 42*µ*m is the separation between electrodes. Interestingly, there is an additional peak in the length distribution of excitatory links at *L* ≈ *d*. This suggests that excitatory signal is transmitted at two spatial scales, one localized scale with transmission limited to nearest nodes situated within several tens of microns and another spatially extended scale with transmission to nodes at millimeters away whereas inhibitory signal is mostly transmitted at the longer millimeter range. We will see later that strong excitation is mainly transmitted at the localized spatial scale.

**Fig 8.**
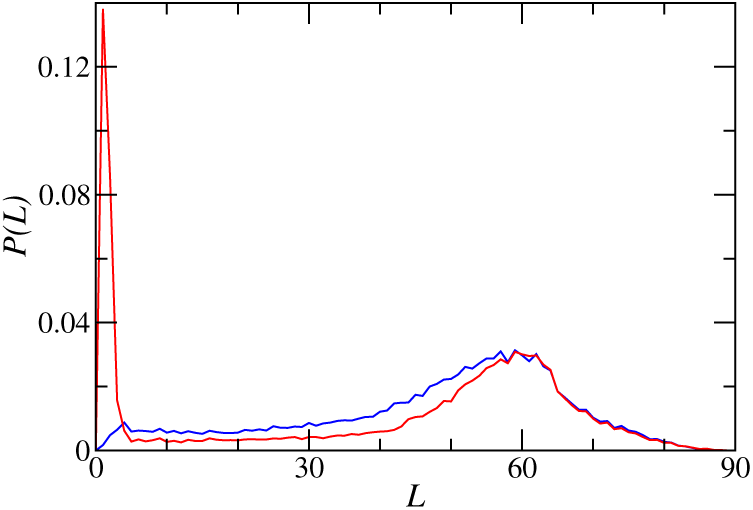
Distribution *P* (*L*) of the physical length *L* of the excitatory (red) and inhibitory (blue) links for DIV25 with *L* measured in units of *d* = 42*µ*m.

### Distributions of incoming and outgoing synaptic strength

We define the average incoming synaptic strength *s*_*in*_(*i*) and the average outgoing synaptic strength *s*_*out*_(*i*) for every node *i* by

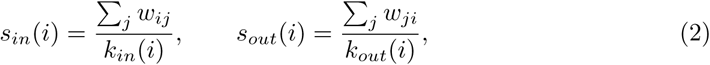

For excitatory nodes *s*_*out*_ > 0 whereas for inhibitory nodes *s*_*out*_ < 0. For nodes with no detectable outgoing links, *s*_*out*_ = 0. For each node, regardless of whether it is excitatory, inhibitory or has no detectable links, *s*_*in*_ can take either sign. Synaptic weights are difficult to be inferred from MEA recordings and results on the distributions of *s*_*in*_ and *s*_*out*_ are generally lacking in previous studies.

As shown in Fig 9, the distribution of *s*_*out*_ is different for excitatory and inhibitory nodes, and for excitatory nodes, the distribution is further different for nodes with positive and negative average incoming synaptic strength *s*_*in*_. For excitatory nodes with *s*_*in*_ > 0, *s*_*out*_ has an approximately lognormal distribution. This result is in accord with the general findings of lognormal distribution of synaptic weights in both in vitro and in vivo studies [28]. For excitatory nodes with *s*_*in*_ < 0 and for inhibitory nodes, the distributions of |*s*_*out*_| have similar functional forms that deviate from lognormal but are better approximated by slightly asymmetric log-exponential distributions.

**Fig 9.**
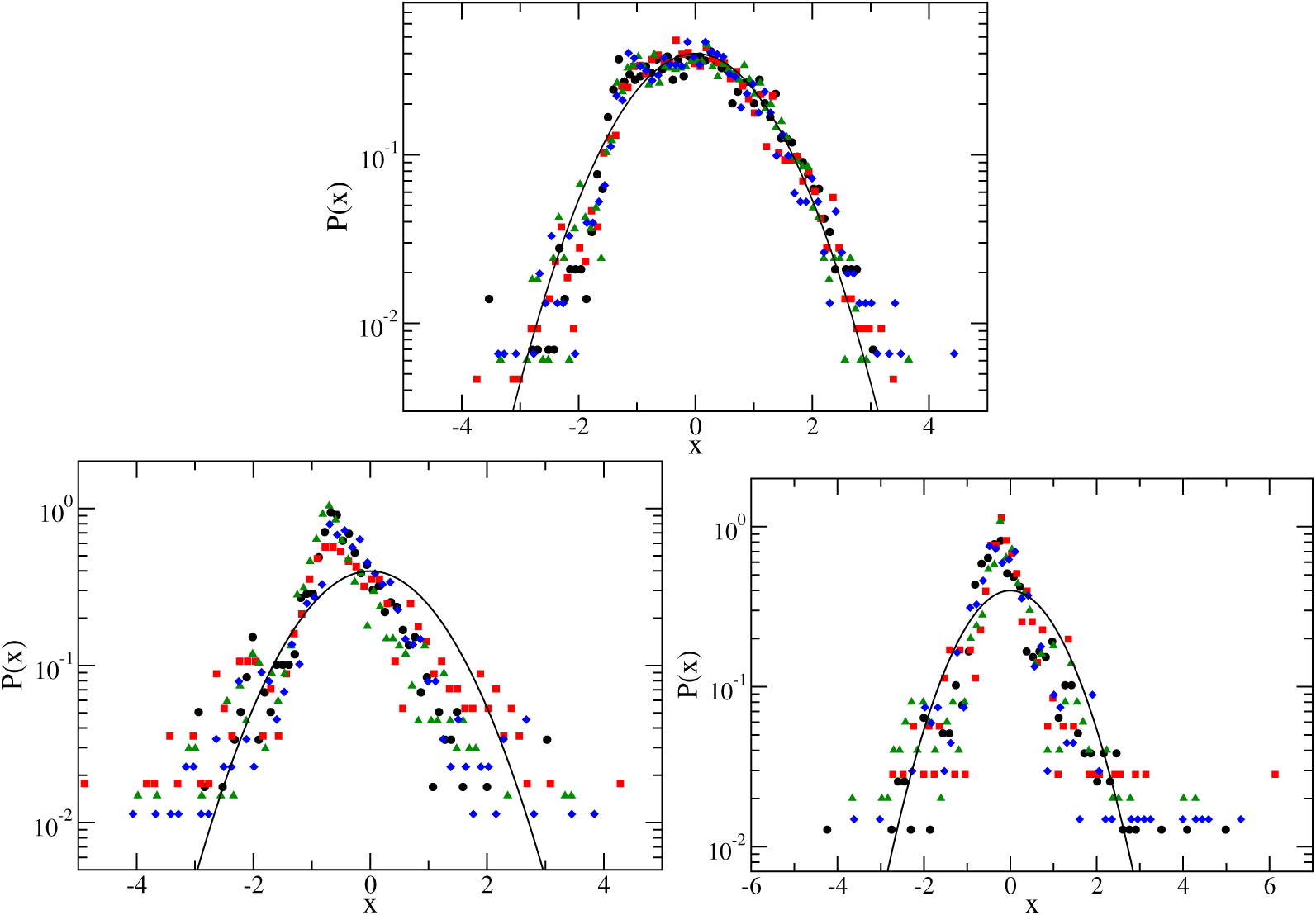
Top panel: Distributions *P* (*x*) of *x* = [ln(*s*_*out*_) − *µ*]*/σ* for excitatory nodes with *s*_*in*_ > 0. Bottom panel: Distributions *P* (*x*) of *x* = [ln |*s*_*out*_| − *µ*]*/σ* for (left panel) excitatory nodes with *s*_*in*_ < 0 and for (right panel) inhibitory nodes. *µ* and *σ* are the mean and standard deviation of ln(*s*_*out*_) respectively. DIV11 (black circles), DIV33 (red squares), DIV52 (green triangles), and DIV66 (blue diamonds) and the solid line is a standardized Gaussian distribution with zero mean and unit variance.

The standardized distributions of |*s*_*in*_| have a universal shape for all DIVs, regardless of whether *s*_*in*_ is greater or less than zero (see Fig 10). These distributions are not lognormal but are better approximated by a highly asymmetric log-exponential distribution. Thus the distributions of both the average incoming and outgoing synaptic strength are non-Gaussian and skewed with long tails. This shows that a small fraction of the nodes have dominantly strong synaptic strength and that the synaptic strength cannot be represented by the mean values.

**Fig 10.**
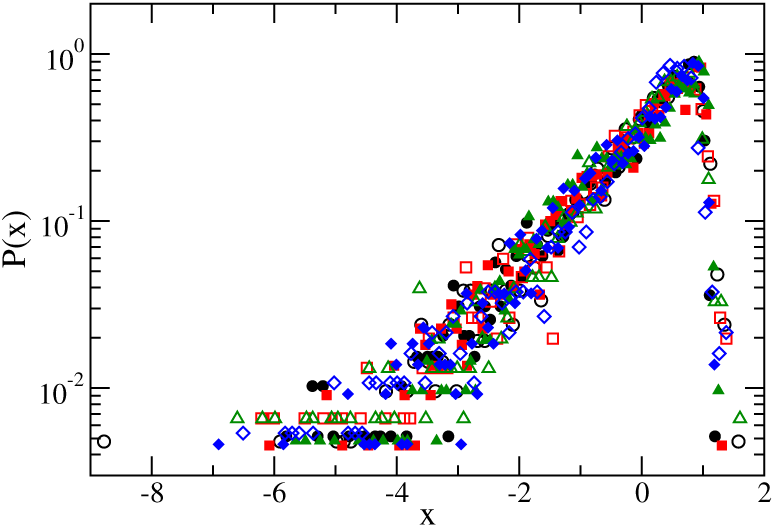
Distributions *P* (*x*) of *x* = [ln(|*s*_*in*_|) − *µ*]*/σ* for nodes with *s*_*in*_ < 0 (open symbols) and *s*_*in*_ > 0 (solid symbols) for DIV11 (black circles), DIV33 (red squares), DIV52 (green triangles), and DIV66 (blue diamonds). *µ* and *σ* are the mean and standard deviation of ln |*s*_*in*_| respectively.

We show the excitatory and inhibitory links of the reconstructed neuronal network for DIV25 in Fig 11. For clarity, we show only the strongest links. We calculated the mean and standard deviation of ln |*w*_*ij*_| separately for excitatory and inhibitory links and show only those links with ln |*w*_*ij*_| being at least the mean value plus 2.5 times the standard deviation. We note the interesting result that the strongest excitatory links mostly have length of *L* ≈ *d*, much shorter than the strongest inhibitory links which mostly have *L* of several tens of *d* of the order of mm. This indicates that strong excitatory signals are mostly transmitted at the localized scale while strong inhibition is mostly long range. Moreover, the strongest inhibitory links are dominated by links from two hubs and the strongest links, excitatory or inhibitory alike, are mainly towards excitatory nodes. We further checked that the shortest links of *L* = *d* are mostly excitatory links towards excitatory nodes and have larger synaptic weight on average.

**Fig 11.**
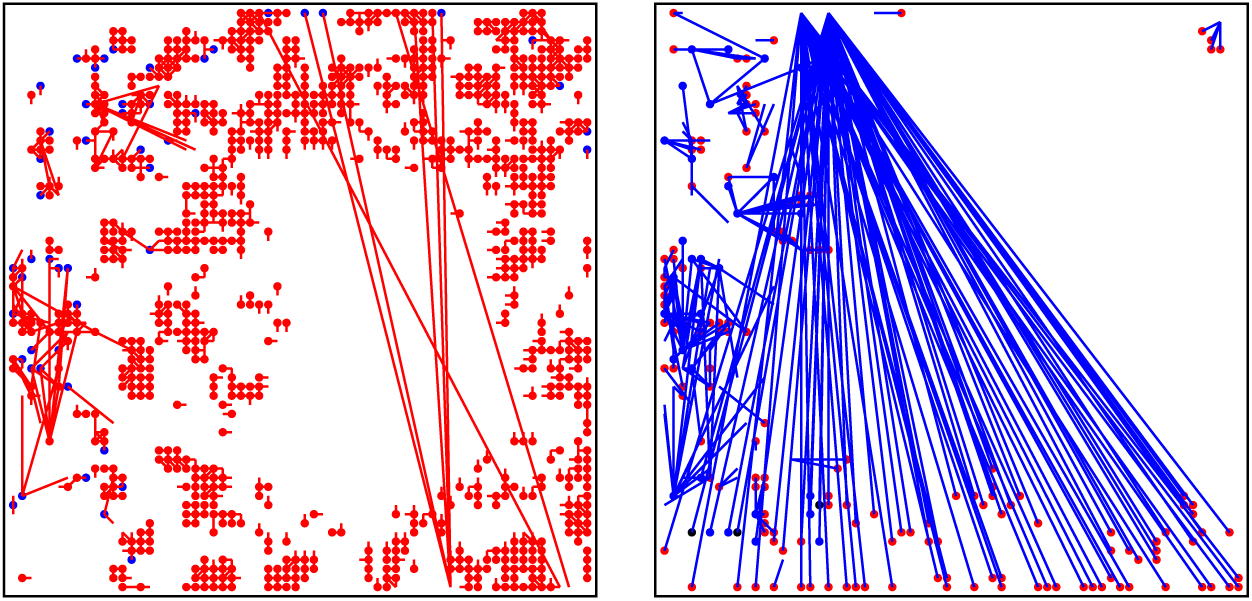
Left panel: The strongest excitatory (left panel) and inhibitory links (right panel) of the reconstructed neuronal network for DIV25 with ln |*w*_*ij*_| ≥ mean + 2.5 standard deviation, with the mean value and standard deviation of ln |*w*_*ij*_| calculated separately for excitatory and inhibitory links. Each excitatory (red) link points from an excitatory node to either an excitatory (red) or inhibitory node (blue). Similarly, each inhibitory (blue) link points from an inhibitory node to either an excitatory (red), inhibitory (blue) or node with no detectable outgoing links (black).

### Comparison of the partial network reconstructed using only measurements from a subset of electrodes and the corresponding subnetwork

As the working area of the MEA probe is more than 4 times the active electrode area, many cells were outside the active electrode area. These cells could form synaptic connections with cells within the electrode area but their spontaneous activities were not recorded. Thus there are missing or hidden nodes in the reconstructed neuronal networks. For bidirectional networks, it has been shown that the presence of hidden nodes that are randomly missed out has no adverse effects on the reconstruction of the network of the measured nodes [29]. The effects of hidden nodes on the reconstruction of directed effective connectivity for general directed networks have yet to be worked out. Thus we studied the effects of hidden nodes in the present problem by reconstructing a partial network using recordings from a subset of 45 by 45 electrodes in the central region of the active electrode area and compared this partial network with the subnetwork of the corresponding 2025 nodes of the reconstructed network using recordings from all 4095 electrodes. We found that the partial network captures correctly 99.8% of the non-existent links and 83.0% of the links in the subnetwork. We compare the spatial distribution of the excitatory, inhibitory nodes and nodes with no detectable links in the subnetwork (which is the same as the central region of the plots shown in Fig 3) and the partial network in Fig 12. There are differences but a great resemblance can be seen.

**Fig 12.**
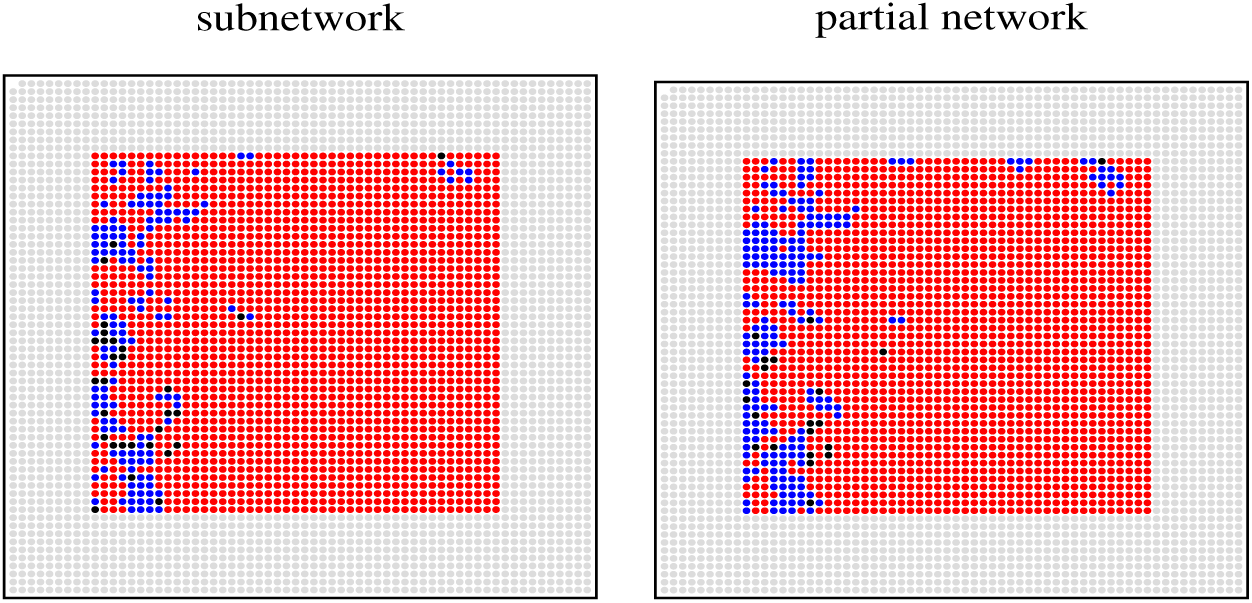
Spatial distribution of excitatory nodes (red), inhibitory nodes (blue) or nodes with no detectable outgoing links (black) for DIV25 in the subnetqwork of the central 45 × 45 electrodes (left) and the partial network reconstructed by using only measurements of a subset of these 2025 electrodes (right).

We further compare the strongest excitatory and inhibitory links of the partial network and the subnetwork using the same criterion defined for strongest links in Fig 11. As can be seen in Fig 13, the partial network can capture the strongest excitatory links in the subnetwork very well. Though there is a larger error for the partial network in obtaining the strongest inhibitory links, the partial network manages to give a correct qualitative picture of the spatial distribution of these strongest inhibitory links. These results hence support that the directed effective connectivity among the measured nodes in the reconstructed networks is not much affected by the missing signal or hidden nodes due to the neural cells lying outside the active electrode area of the MEA probe.

**Fig 13.**
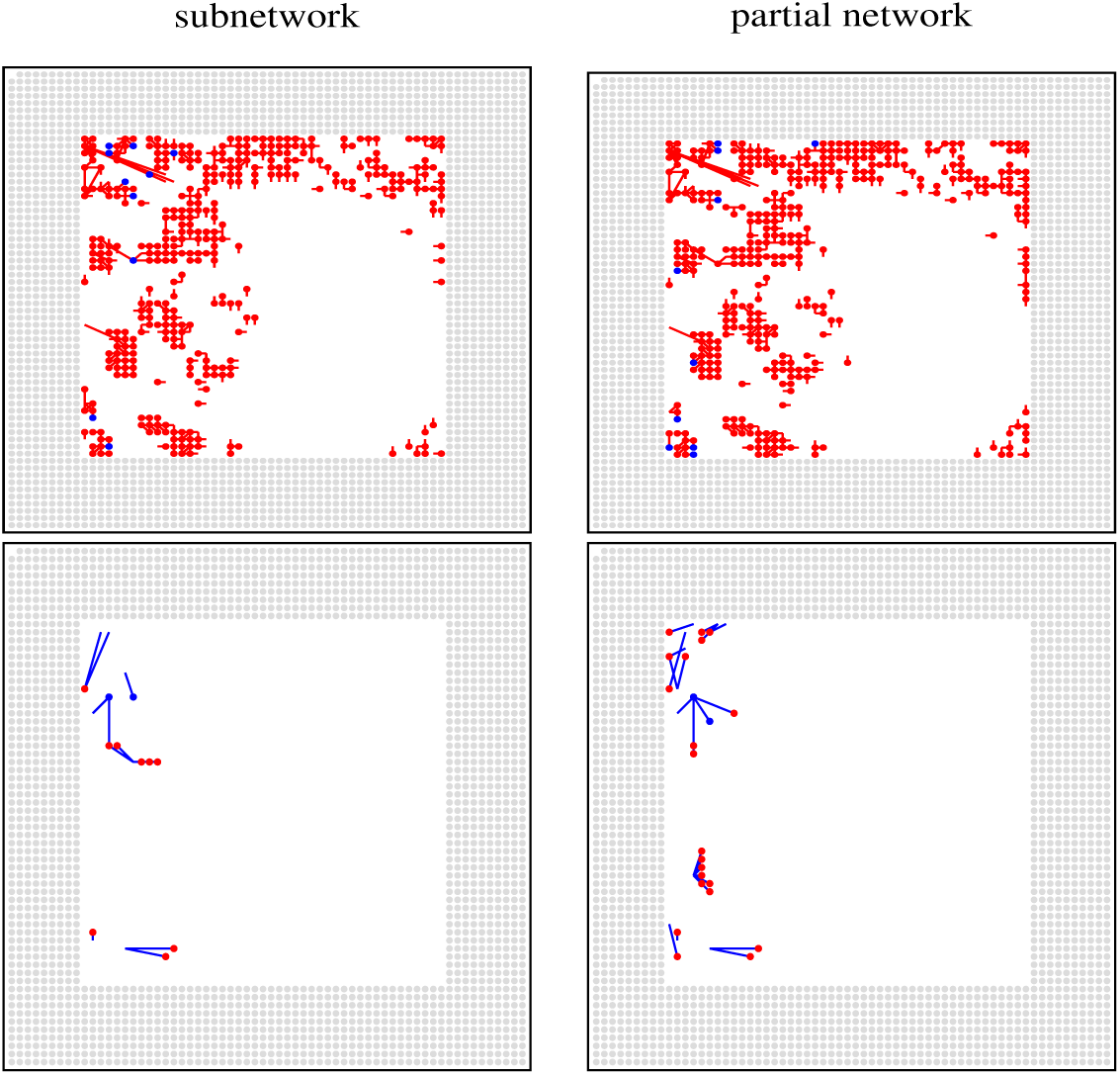
Top panel: The strongest excitatory links of the subnetwork (left) and the partial network (right) for DIV25. Bottom panel: The strongest excitatory links of the subnetwork (left) and the partial network (right) for DIV25.

## Discussion

Revealing directed connectivity of neuronal networks from electrophysiological data recorded by large-scale MEA is a challenging inverse problem. Existing methods focus on functional connectivity which measures statistical dependencies of the activities of neurons. To address questions regarding connectivity and dynamics and network structure and functions of neuronal networks, the relevant quantity should, however, be effective connectivity which measures direct causal interactions and not functional connectivity. As it is natural to infer effective connectivity using a specific dynamical model, it is commonly thought that effective connectivity is model-dependent. This further adds to the difficulty to generally investigate directed effective connectivity of neuronal networks. One established technique for estimating functional connectivity makes use of the cross-correlation of spiking activity of neurons. It was demonstrated theoretically that for a generic class of models and not just for a specific model, the effective connectivity matrix is contained in a theoretical relation between the time-lagged cross-covariance and the equal-time cross-covariance [13]. Making use of this relation, a method has been developed to recover effective connectivity [13] and this method was validated for models with different dynamics and coupling using numerical data [13, 16]. This covariance-relation based method involves a calculation of a principal matrix logarithm, which is very sensitive to noise in the data. When we applied it directly to the MEA recordings, we obtained an unphysical complex matrix for the supposed effective connectivity matrix. A complex matrix was also found when this covariance relation was used directly to estimate directed effective connectivity of a cortical network of 68 regions from fMRI recordings and this motivated the development of a Lyapunov optimization procedure [11]. Here, we showed that the unphysical complex matrix problem can be avoided by first applying a moving average filter to the MEA recordings to reduce random noise (c.f. Materials and methods for the method of reconstruction). Moreover, we extended the originally proposed clustering analysis using a Gaussian mixture model [13] to include also a single-Gaussian fit with outliers. We made additional assumptions that the neuronal networks are sparse and outgoing links of any node, if exist, have to be all excitatory or all inhibitory. We have shown that this new covariance-based method, the original proposed method together with the modifications, can estimate the directed and weighted effective connectivity matrix of neuronal networks from recordings taken by MEA of over 4000 electrodes. Specifically, the simulated spiking activity using the estimated directed and weighted effective connectivity in a stochastic LIF model can capture the spontaneous network bursts of the measured spiking activity and retrieve the null-nodes in the time-window of observation reasonably well.

The balance between excitation and inhibition in the cortex is believed to play an important role in executing proper brain functions and disruption of such a balance may underlie the behavioural deficits that are observed in conditions such as autism and schizophrenia [30–34]. To study and gain insights of excitation/inhibition balance in neuronal networks, it is important that the methods estimating connectivity can detect excitatory and inhibitory links equally well. The cross-covariance based method for functional connectivity has a considerably weaker sensitivity for detecting inhibition and a recently introduced filtering approach [10] overcomes this problem but the detected excitation/inhibition ratio from MEA of 4096 electrodes tends to stabilize only when the recording time is of of the order of 30 min. Our method can detect excitatory and inhibitory links equally well and we successfully detected both excitatory and inhibitory links from the same type of MEA using only a recording time of 5 min and identified nodes of the neuronal networks as excitatory, inhibitory or with no detectable outgoing links. The fraction *f*_*I*_ of inhibitory nodes is 0.13-0.31 for the 8 DIVs studied, which overlaps with the range 0.15-0.30 of measured fractions of inhibitory neurons in various cortical regions of monkey [25]. In this sense, the reconstructed neuronal networks are excitation/inhibition balanced as found in monkey cortex.

Neuronal networks are expected to be organized and should thus be highly nonrandom in order to transmit signals efficiently and to carry out their functions effectively. Similar to the chemical synapse network of C. elegans [20], the reconstructed neuronal networks have small-world topology, and it has been shown that small-word network allows efficient transmission of information [35]. The nonrandom feature of local cortical circuits has been well documented by an over-representation of bidirectional connected pairs as compared to a random network of the same connection probability [21–24]. In the reconstructed neuronal networks, the number of bidirectionally connected pairs was indeed found to exceed at least four times the expected number for the random network. Bidirectional connected pairs can play a direct role in forming feedback control in neuronal circuits. We estimated the functional connectivity for DIV25, DIV45 and DIV66 by using the cross-covariance based methods [10] and bidirectionally connected pairs are not found. Neuronal networks are also expected to be robust against random failure or targeted attacks. There were studies in network science indicating that optimization of the network topology of undirected networks against both random failures and attacks can be achieved by a bimodal degree distribution [26, 27]. Interestingly, the incoming degree distribution of the reconstructed neuronal networks is found to be bimodal. It is thus interesting to investigate whether a bimodal incoming degree distribution would optimize the robustness of directed networks against certain attacks.

The incoming and outgoing degrees of the neuronal networks have distinctive distributions. The outgoing degree distribution for both excitatory and inhibitory nodes has a long tail. The maximum incoming degree is of the order of 100 while the maximum outgoing degree is of the order of 1000. Thus hubs are feeders rather than receptors and there are both excitatory and inhibitory feeders locating mainly on the top and bottom rows of the MEA probe. The excitatory and inhibitory incoming degrees have bimodal distributions and nodes in the two modes are spatially segregated. The nodes in the modes of larger 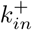 and larger 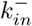 occupy the upper-quarter and lower-quarter regions of the MEA probe respectively showing clearly that nodes do not exhibit both large excitatory and large inhibitory incoming degrees. These results suggest some form of differentiation of roles of the nodes in the neuronal circuits in receiving excitatory and inhibitory signals. The distribution of the physical length of excitatory links has two peaks indicating that excitatory signals are transmitted at two distinct spatial scales, one highly localized scale with transmission to nearby nodes within several tens of microns and another spatially extended scale with transmission to nodes at millimeters away. The distribution of the physical length of inhibitory links has only one broad peak at the spatially extended scale. We further found that strong excitatory signals are mostly transmitted locally to nearby excitatory nodes while strong inhibitory signals are mostly transmitted at the spatially extended millimeter scale from a few hubs. Long range inhibitory projections from GABAergic neurons have been demonstrated in adult rat hippocampus [36, 37] and the hub function of these neurons has been suggested [38, 39].

Incoming synaptic strength is difficult to measure directly and results on their distributions have been lacking in the literature. The present covariance-relation based method allows us to estimate the synaptic weights of all the links and, as a result, we can calculate the distributions of incoming and outgoing synaptic strength of the nodes. We found that these distributions are described by lognormal or log-exponential functional form (Fig 9 and 10) and they are in line with the ubiquitous lognormal-like distributions of many physiological and anatomical features at different levels in the brain [28]. The significance of these skewed non-Gaussian distributions with long tails is that a small fraction of nodes have dominantly strong synaptic strength suggesting the possibility that the bulk of the information flow occurring mostly through these nodes [28]. Understanding how this lognormal-like synaptic strength might be related to the spiking dynamics or the general relation between connectivity and dynamics will be a topic of great interest for future studies.

## Materials and methods

### Ethics statement

Cortices from embryonic day 17 (E17) Wistar rat were used in the experiments. The experimental protocol was evaluated and approved by the Institutional Animal Care and Use Committee of Academia Sinica (AS IACUC) with Protocol ID: 12-12-475.

### Neuronal cultures and experimental set-up

Tissues were dissected from 2 to 3 rats and digested with 0.125 % trypsin for 15 min at 37°C. A small drop (100 *µ*l) of cell suspension was plated on the working area of the complementary metal-oxide-semiconductor (CMOS)-based high density multielectrode array (HD-MEA), which was pre-coated with 0.1 % Poly-D-lysine (Sigma P6407) and 0.1 % adhesion proteins laminin (Sigma L2020) (see Fig 14). A total of about 6 × 10^4^ cells were plated on a working area of about 6 mm × 6mm of the HD-MEA chip resulting on a cell density of about 1670 cells/mm^2^. After plating on the chip, cultures were filled with 1 ml of culture medium [DMEM (Gibco 10569) + 5% FBS (Gibco 26140) + 5% HS (Gibco 16050) + 1% PS (Gibco 15140)] and placed in a humidified incubator (5% CO_2_, 37°C). Half of the medium was replaced by Neurobasal medium supplemented with B27 [Neurobasel medium (Gibco. 21103) + 2% 50X B27 supplement (Gibco. 17504) + 200 *µM* GlutaMAX (Gibco 35050)] twice a week.

**Fig 14.**
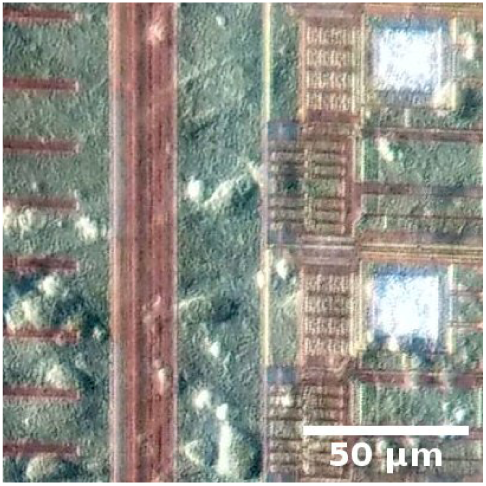
Surface of high-density CMOS MEA. The line shows a scale of 50*µ*m.

The HD MEA probe (HD-MEA Arena, 3Brain AG) has 4096 electrodes arranged in a 64 by 64 square grid. The size of each square electrode is 21 *µ*m by 21*µ*m and the electrode pitch is 42 *µ*m which gives an active electrode area of 2.67 mm by 2.67 mm. Neuronal activities from HD MEA were recorded with the recording device (BioCAM, 3Brain AG) and the associate software (BrainWave 2.0, 3Brain AG) at 7.06 kHz. One electrode was used for calibration purpose so there were 4095 electrodes that recorded signals. Spontaneous activities were recorded from 6 to over 60 days in vitro. Samples were placed into the recording device 10 min before the recording in order to prevent the effects of vibration. Each experimental session lasted 5 min and recorded in dark since the CMOS is a light active material.

### Method of reconstruction

We assumed that the activity *x*_*i*_(*t*), recorded by each of the electrodes, is stationary and considered a generic dynamical model in which the time evolution of the fluctuations around the average value, *δx*_*i*_(*t*) = *x*_*i*_(*t*) − ⟨*x*_*i*_(*t*)⟩, is governed by a system of linear stochastic differential equations

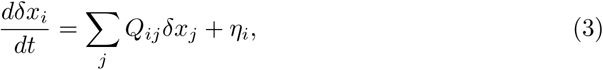

where ⟨…⟩ denotes a time average, *Q*_*ij*_ = *w*_*ij*_(1 − *δ*_*ij*_) + *Q*_*ii*_*δ*_*ij*_ are the elements of the interaction matrix **Q** and *η*_*i*_ is a Gaussian white noise of zero mean with

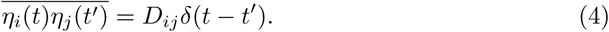

When the activity *x*_*j*_ affects the activity *x*_*i*_, there is a link from node *j* to node *i* with synaptic weight *w*_*ij*_ otherwise *w*_*ij*_ = 0. Thus *w*_*ij*_ give the directed and weighted effective connectivity of the neuronal network. We note that such a linear stochastic system of stationary fluctuating activity can arise even for systems whose activity obey nonlinear dynamics [13]. For such a model, the time-lagged covariance matrix **K**(*τ*) and the equal-time covariance matrix **K**(0) [11, 13, 40, 41] of the measurements are related by the interaction matrix **Q**:

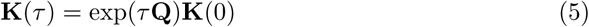

The elements of **K**(*τ*) and **K**(0) are defined by

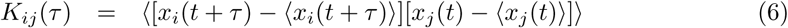

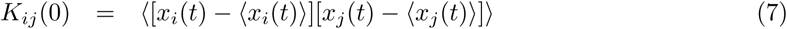

respectively. This relation implies that the off-diagonal elements of **M** = log(**K**(*τ*)**K**(0)^−1^)*/τ*, where log is the principal matrix logarithm, would separate into two groups, one corresponds to *w*_*ij*_ = 0 and another to *w*_*ij*_ ≠ 0. It has been shown [13] that by clustering analysis of *M*_*ij*_ using a Gaussian mixture model, *w*_*ij*_ can be inferred. We extended this proposed clustering analysis to include also a single-Gaussian fit with outliers.

For the reconstruction, we only made use of measurements *y*_*i*_(*t*), *i* = 1, 2, …, 4095 taken during which all the 4095 electrodes were recording properly. The principal matrix logarithm is very sensitive to noise in the measurements and a calculation of **K**_*y*_(*τ*) and **K**_*y*_(0) directly from these MEA recordings *y*_*i*_(*t*) [as defined in Eqs (6) and (7) with *x*_*i*_ replaced by *y*_*i*_] would lead to the matrix log(**K**_*y*_(*τ*)**K**_*y*_(0)^−1^) having complex elements. This has also been found for fMRI measurements [11]. We avoided this problem by applying a moving average filter to the MEA recordings to reduce random noise. The moving average filter is a simple digital low-pass filter that is optimum for reducing random noise while retaining sharp transitions in the data [43]. Specifically, we took *x*_*i*_(*t*) = [*y*_*i*_(*t*) + *y*_*i*_(*t* + Δ)]*/*2, where Δ is the sampling time interval for all the valid MEA recordings *y*_*i*_(*t*), *i* = 1, 2, …, 4095, and calculated **K**(*τ*) and **K**(0) from *x*_*i*_(*t*) with *τ* = Δ. The resulting matrix **M** was found to be real. Then we adopted the clustering method of Ref. [13] with several modifications to cluster *M*_*ij*_’s for each fixed *j* into two groups, one corresponds to *w*_*ij*_ = 0 and the other to *w*_*ij*_ ≠ 0.

We assumed that the outgoing links of each node, if exist, can only be all excitatory with *w*_*ij*_ > 0 or all inhibitory with *w*_*ij*_ < 0. This is in accord to neurons are either excitatory or inhibitory. We first fitted the values of *M*_*ij*_’s with fixed *j* by a Gaussian mixture model of two components, namely by a sum of two Gaussian distributions, using MATLAB ‘fitgmdist’, which return the means, *µ*_1_ and *µ*_2_, and standard deviations, *σ*_1_ and *σ*_2_, of the two Gaussian distributions, and the relative proportion of the two components. We divided the results into three cases according to the separation between the two fitted Gaussian distributions: (i) the two distributions are well separated with |*µ*_1_ − *µ*_2_| > *σ*_1_ + *σ*_2_, (ii) the two distributions are very close to one another with |*µ*_1_ − *µ*_2_| < max(*σ*_1_, *σ*_2_), and (iii) the in-between case with max(*σ*_1_, *σ*_2_) ≤ |*µ*_1_ − *µ*_2_| ≤ *σ*_1_ + *σ*_2_. For case (i), we continued with this two-Gaussian fit. For case (ii), we fitted the data again by a single Gaussian distribution instead. For case (iii), we chose either the two-Gaussian fit or the single-Gaussian fit according to the Bayesian information criterion (BIC) for fitting models selection [42, 44].

When using the two-Gaussian fit, we thus inferred the unconnected component for *w*_*ij*_ = 0 as the component of a relative proportion greater than 0.6 based on the assumption that the neuronal cultures are sparse networks or as the component whose mean is closer to zero when none of the relative proportions exceeds 0.6. We obtained the posterior probability *p* of each data point belonging to the unconnected component using MATLAB ‘cluster’. For each *M*_*ij*_, if *p* > 0.5 we inferred *w*_*ij*_ = 0 otherwise, we inferred *w*_*ij*_ to be given by *M*_*ij*_ − ⟨*M*_*ij*_|*w*_*ij*_ = 0⟩_*i*_ and ⟨*M*_*ij*_|*w*_*ij*_ = 0⟩_*i*_ is the average of *M*_*ij*_ with inferred *w*_*ij*_ = 0 over *i* for fixed *j*. Thus the synaptic weights *w*_*ij*_ of the outgoing links of each node *j* have the same sign. Depending on whether *w*_*ij*_ is greater than or less than zero, the node was inferred to be excitatory or inhibitory (c.f. Fig 15).

**Fig 15.**
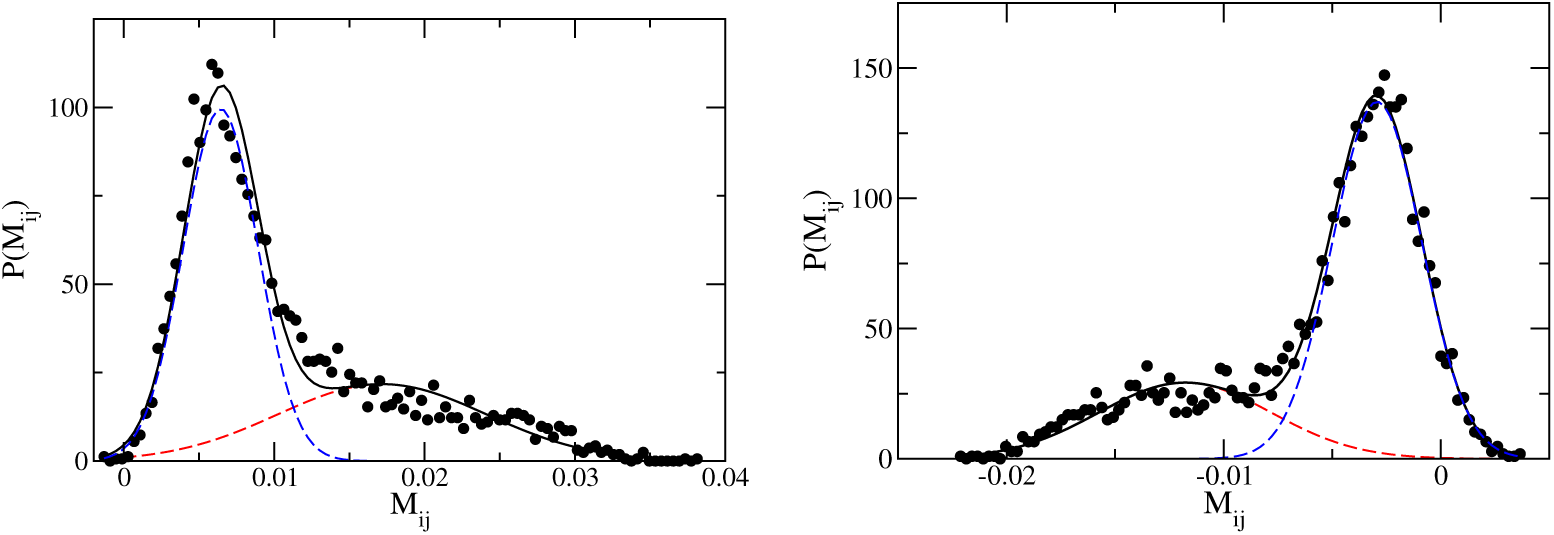
Typical distributions *P* (*M*_*ij*_) of *M*_*ij*_ (circles) with *j* fixed for an excitatory node in DIV66 (left panel) and an inhibitory node in DIV 11 (right panel) inferred by using a Gaussian-mixture model (solid black lines) of two components (blue and red dashed lines).

When using the single-Gaussian fit, we identified the data points that are well fitted by the Gaussian, denoted by *P*_*G*_ of mean *µ*, with *w*_*ij*_ = 0 and the outliers, which are data points deviate significantly from the fitted Gaussian, with *w*_*ij*_ ≠ 0. For this purpose, we let *x*_*E*_ and *x*_*I*_ be

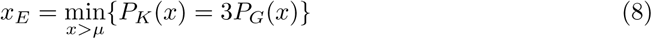

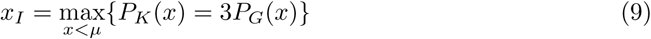

where *P*_*K*_ is the distribution of *M*_*ij*_ obtained by the Kernel density estimation and calculated the number of data points (*N*_1_) satisfying *M*_*ij*_ > *x*_*E*_ and the number of data points (*N*_2_) satisfying *M*_*ij*_ < *x*_*I*_. The outliers are the *N*_1_ *M*_*ij*_’s if *N*_1_ > *N*_2_ or the *N*_2_ *M*_*ij*_’s if *N*_1_ < *N*_2_. The identified outliers were inferred to have *w*_*ij*_ = *M*_*ij*_ − ⟨*M*_*ij*_|*w*_*ij*_⟩ = 0 _*i*_. If both *N*_1_ = *N*_2_ = 0, then node *j* was inferred to have no detectable outgoing links.

## Code availability

The MATLAB program for clustering analysis of the calculated matrix **M** to give the adjacency matrix elements *A*_*ij*_ and coupling *w*_*ij*_ can be found in the Open Science Framework (OSF) https://osf.io/5dc4j/

### Stochastic leaky-integrate-and-fire spiking neuron model

The dynamics of a stochastic leaky-integrate-and-fire (LIF) neuron is governed by the following equation

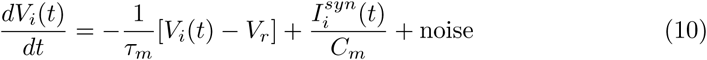

where *V*_*i*_(*t*) is the membrane potential of neuron *i, V*_*r*_ is the resting potential, *C*_*m*_ and *G*_*m*_ are the membrane capacitance and membrane conductance, *τ*_*m*_ = *C*_*m*_*/G*_*m*_ is the decay constant, and 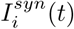 is the synaptic current input from the presynaptic nodes. When *V*_*i*_(*t*) exceeds a threshold *V*_*th*_, node *i* generates an action potential or spike at that instant and *V*_*i*_ is reset to *V*_*r*_. The synaptic current 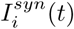 is given by

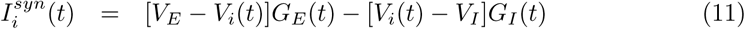

where *V*_*E*_ and *V*_*I*_ are the reversal potentials satisfying *V*_*I*_ ≤ *V*_*r*_ < *V*_*th*_ ≤ *V*_*E*_, and *G*_*E*_(*t*) and *G*_*I*_ (*t*) are the excitatory and inhibitory synaptic conductances given by

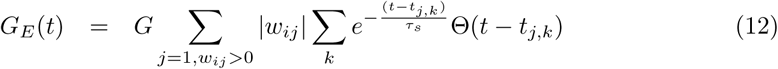

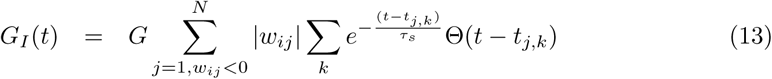

where *G* is a typical synaptic conductance, *w*_*ij*_ are the synaptic weights, *τ*_*s*_ is the decay constant of the synaptic conductances, Θ(*t*) is the Heaviside step function and *t*_*j,k*_ is the spiking time of the *k*-th spike of the presynaptic neuron *j*.

To simplify the notations, we worked with the dimensionless membrane potential, which is defined by

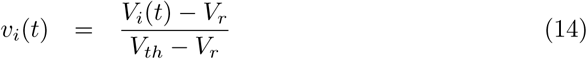

with 0 ≤ *v*_*i*_ ≤ 1 then the equations become

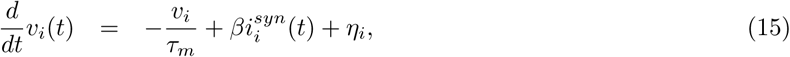

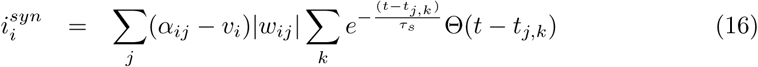

where *β* = *G/C*_*m*_,

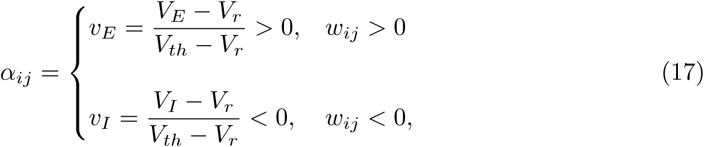

and *η*_*i*_ is a Gaussian white noise of zero mean defined by Eq (4) with *D*_*ij*_ = *σ*^2^. When *v*_*i*_ ≥ 1, *v*_*i*_ is reset to 0.

In the numerical simulations, we used *τ*_*m*_ = 20 ms, *τ*_*s*_ = 2 ms, *v*_*E*_ = 6 and *v*_*I*_ = −2 (which follow from *V*_*r*_ = −60 mV, *V*_*th*_ = −50 mV, *V*_*E*_ = 0 mV and *V*_*I*_ = −80 mV) [45] and *σ* = 2. We took *β* = 1000 and *β* = 40 respectively for the reconstructed networks and the control randomly-shuffled networks with standardized Gaussian synaptic weights. The time step Δ*t* of the simulations was taken to be the sampling time interval Δ*t*=1/7060*s* ≈ 0.142 ms and we obtained the spiking activity of each node *i* using the measured spikes of all its presynaptic nodes (with *w*_*ij*_ ≠ 0) as inputs.

